# Gene interaction network analysis in multiple myeloma detects complex immune dysregulation associated with shorter survival

**DOI:** 10.1101/2023.04.05.535155

**Authors:** Anish K. Simhal, Kylee H. Maclachlan, Rena Elkin, Jiening Zhu, Larry Norton, Joseph O. Deasy, Jung Hun Oh, Saad Z. Usmani, Allen Tannenbaum

**Affiliations:** Department of Medical Physics, Memorial Sloan Kettering Cancer Center, New York, NY; Department of Medicine, Memorial Sloan Kettering Cancer Center, New York, NY; Department of Applied Mathematics & Statistics, Stony Brook University, Stony Brook, NY; Departments of Computer Science and Applied Mathematics & Statistics, Stony Brook University, Stony Brook, NY

**Keywords:** multiple myeloma, network analysis, RNA-sequencing, copy number aberration, immune system, DNA damage.

## Abstract

The plasma cell cancer multiple myeloma (MM) varies significantly in genomic characteristics, response to therapy, and long-term prognosis. To investigate global interactions in MM, we combined a known protein interaction network with a large clinically annotated MM dataset. We hypothesized that an unbiased network analysis method based on large-scale similarities in gene expression, copy number aberration, and protein interactions may provide novel biological insights. Applying a novel measure of network robustness, Ollivier-Ricci Curvature, we examined patterns in the RNA-Seq gene expression and CNA data and how they relate to clinical outcomes. Hierarchical clustering using ORC differentiated high-risk subtypes with low progression free survival. Differential gene expression analysis defined 118 genes with significantly aberrant expression. These genes, while not previously associated with MM, were associated with DNA repair, apoptosis, and the immune system. Univariate analysis identified 8/118 to be prognostic genes; all associated with the immune system. A network topology analysis identified both hub and bridge genes which connect known genes of biological significance of MM. Taken together, gene interaction network analysis in MM uses a novel method of global assessment to demonstrate complex immune dysregulation associated with shorter survival.

**STATEMENT OF SIGNIFICANCE:** Multiple myeloma has heterogenous clinical outcomes which are not well predicted by current prognostic scoring systems. Global assessment of gene-protein interactions using Ollivier-Ricci Curvature produces clusters of patients with defined prognostic significance, with high-risk groups harboring complex gene dysregulation impacting immune function.

## INTRODUCTION

The plasma cell cancer multiple myeloma (MM) has highly heterogenous clinical outcomes, with a key determinant of response to treatment being genomic driver events. The most common recurrent genomic events are hyperdiploidy, with a predominance of gains in chromosomes 3, 5, 7, 9, 11, 15, 19, and 21, and canonical chromosomal translocations affecting the immunoglobulin heavy chain on chromosome 14 (1). MM harbors relatively few recurrent point mutations compared with many other cancers, with only *NRAS, KRAS, TP53, FAM46C* and *DIS3* having a prevalence above 10% (2).

Prognostic scoring updates have expanded the International Staging System (ISS) to incorporate several chromosomal translocations [t(4;14), t(14;16)] and copy number aberrations (CNA; deletion17p, gain/amplification1q), with each feature being considered as an individual event (3,4). It has been recognized, however, that neither these features nor somatic mutations are sufficient to define prognosis, with more extensive genomic assessments required to accurately predict biological behavior.

Previous studies have described various genomic subtypes of MM using RNA-sequencing (RNA-Seq) and/or CNA data (5–10). The subtypes identified by these methods tend to be dominated by a single genomic event (i.e., hyperdiploidy, t(11;14), t(4;14), high proliferation index) or a combination of previously described events (i.e., complex hyperdiploidy with gain1q and monosomy 13) (9).

Here, we consider that integrating data from a comprehensive systems view, incorporating complex interactions between multiple genes in a network, may define patterns of biological behavior not captured by individual genomic events. Recently, a novel measure of network robustness, Ollivier Ricci curvature (ORC), has been used to characterize breast and ovarian cancers (11,12) and other pathological states (13). ORC measures the ability of a given connection or interaction, between a pair of nodes — here being genes — to withstand perturbation, considering both local and global connectivity in assessing the robustness of each pathway (see **Methods** for a detailed description). In the context of cancer genomics, positive curvature implies that there are multiple, robust active pathways for communication between the two genes. This edge, or connection, can be described as “hub-like”. Negative curvature implies that if the connection between two genes is impacted, the effect is relatively greater because of lack of direct feedback controls; this edge can be considered “bridge-like”. Therefore, ORC analysis predicts the effect of changes in gene expression within a wider network as opposed to just the individual gene.

We utilize the ongoing Multiple Myeloma Research Foundation (MMRF) multi-site longitudinal clinical registry study, which follows patients newly diagnosed with MM and collects both clinical and genomic information periodically (9,14). The project, entitled CoMMpass (Relating Clinical Outcomes in Multiple Myeloma to Personal Assessment of Genetic Profile), has over a thousand patients enrolled in the latest interim analysis (IA19), and represents the largest publicly available MM genomic data repository. The dataset includes clinical information, RNA sequencing (RNA-Seq) information, copy number aberration (CNA), among others. To understand the relationship between genes, we used a gene interactome derived from the Human Protein Reference Database (HPRD) (15).

In this study, we apply an innovative geometric network analysis that integrates complex gene-product interactions to characterize global patterns of MM biological behavior. Hierarchical clustering defined groups of patients having different survival times, despite similar ISS distributions. We identified 118 genes having significantly aberrant expression, most of which are previously unassociated with MM, and 8 genes with prognostic capabilities which are part of the immune system. These genes are not just hub genes, but bridge genes which help modulate connections between two larger hub genes. Here, we demonstrate that protein-gene interaction network analysis in MM demonstrates complex immune dysregulation which associates with shorter survival.

## METHODS

In this study, we perform a comprehensive geometric network analysis that integrates complex gene-product interactions to characterize patterns in biological states. The methodology is mathematically well-defined and has no fitting parameters, with an outline of the process illustrated in **Figure 1**.

**Figure 1.**
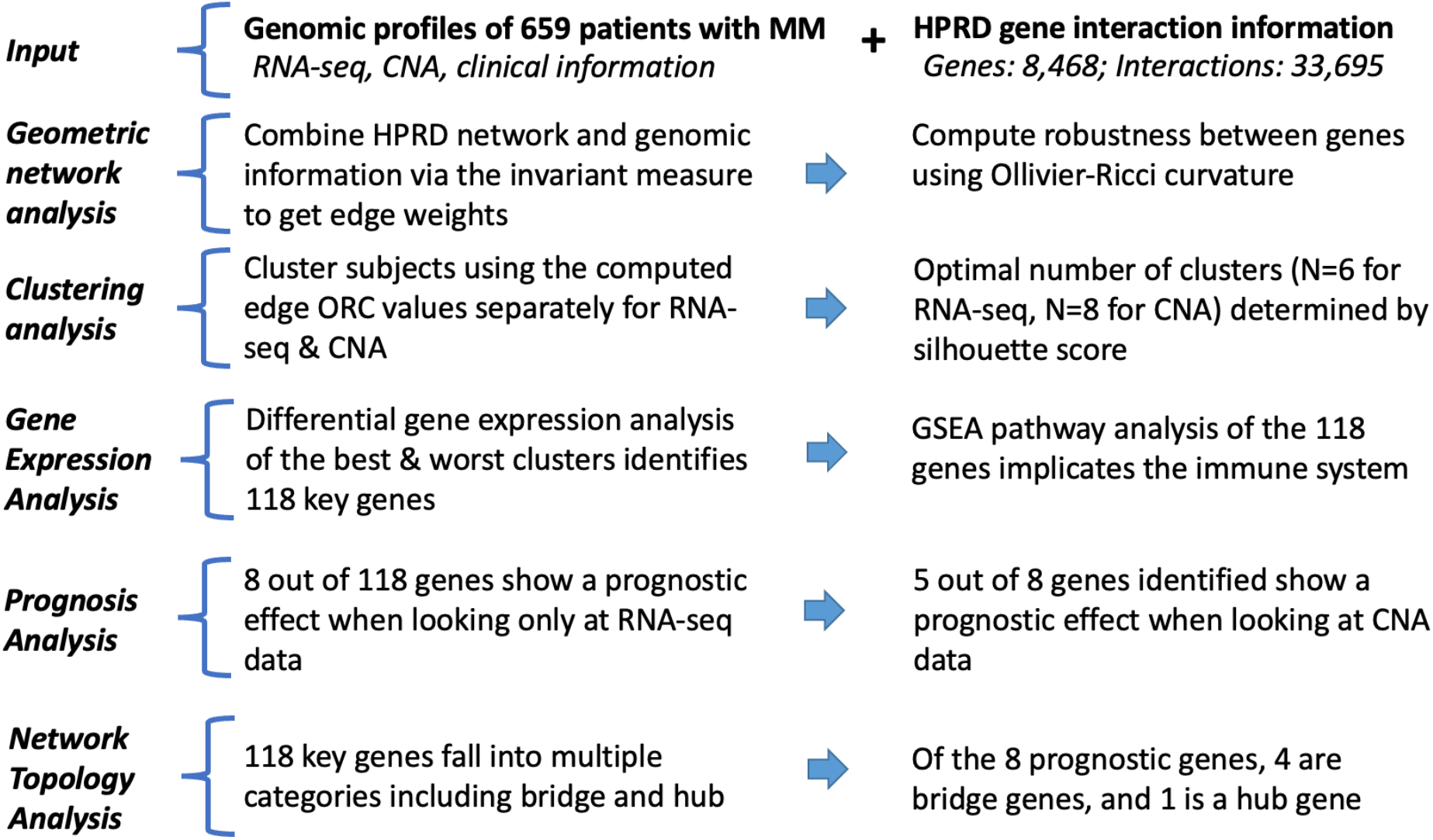
Overview of the data processing pipeline. This study uses a novel measure of network robustness, Ollivier-Ricci curvature, to examine genes associated with shorter progression free survival in multiple myeloma. RNA-Seq: RNA-sequencing; HPRD: Human Protein Reference Database; CNA: copy number aberration; ORC: Ollivier-Ricci curvature; GSEA: gene set enrichment analysis.

### Genomic data

The MMRF CoMMpass dataset (release iteration: IA19), available to all researchers at www.research.mmrf.org, includes clinical information, RNA-Seq gene expression, and CNA data collected over time. Further information on the data collection and curation methods has previously been published (9). For inclusion in this study, subjects must have RNA-Seq and CNA data extracted from the bone marrow prior to the start of treatment and both demographic and survival information. For the RNA-Seq data, the data provided by the MMRF was preprocessed using the SALMON toolbox (16), included filtering unstranded immunoglobulin values, and was normalized as transcripts per million (TPM) and log-transformed. For the CNA data, the data provided by the MMRF was preprocessed using GATK (9).

Hyperdiploidy defined by more than 2 gains involving >60% of the chromosome affecting chromosomes 3, 5, 7, 9, 11, 19 or 21. Mutational signatures were assessed using *mmsig* (https://github.com/UM-Myeloma-Genomics/mmsig), a fitting algorithm designed specifically for MM, to estimate the contribution of each mutational signature in each sample. APOBEC-mutational activity was calculated by combining SBS2 and SBS13, with the top 10% being defined as hyper-APOBEC (https://cancer.sanger.ac.uk/signatures/sbs/). The complex structural variant chromothripsis was defined by manual curation according to previously published criteria (17).

### Gene-product interaction data

For network analysis on gene-product interactions, we used the curated network given by the Human Protein Reference Database (HPRD) (15). The database consists of 9,600 genes and notates 36,822 interaction pairs. We used the largest connected component of shared information among the HPRD, RNA-Seq, and CNA data sets, which included 8427 of 9600 potential genes.

### Ollivier Ricci curvature

ORC integrates both local and global connectivity in assessing the robustness of each interaction as characterized by the numerous feedback loops in a network modeled by a weighted graph or Markov chain (18). Robustness, in this context, is defined as the ability of a system to return to its original state following a perturbation. The ORC calculation is based on the ratio of an intrinsic graph distance, capturing the metric properties of the network, to a distance defined via optimal transport theory between the distributions of neighboring genomic values connected to a given node. Capturing the sample-dependent pattern of curvature weighted edges provides a powerful network-wide signature that integrates non-local information; illustrated in **Figure 2**, examples zero, positive and negative curvature. ORC was calculated as per previous descriptions (11) and is defined below.

**Figure 2.**
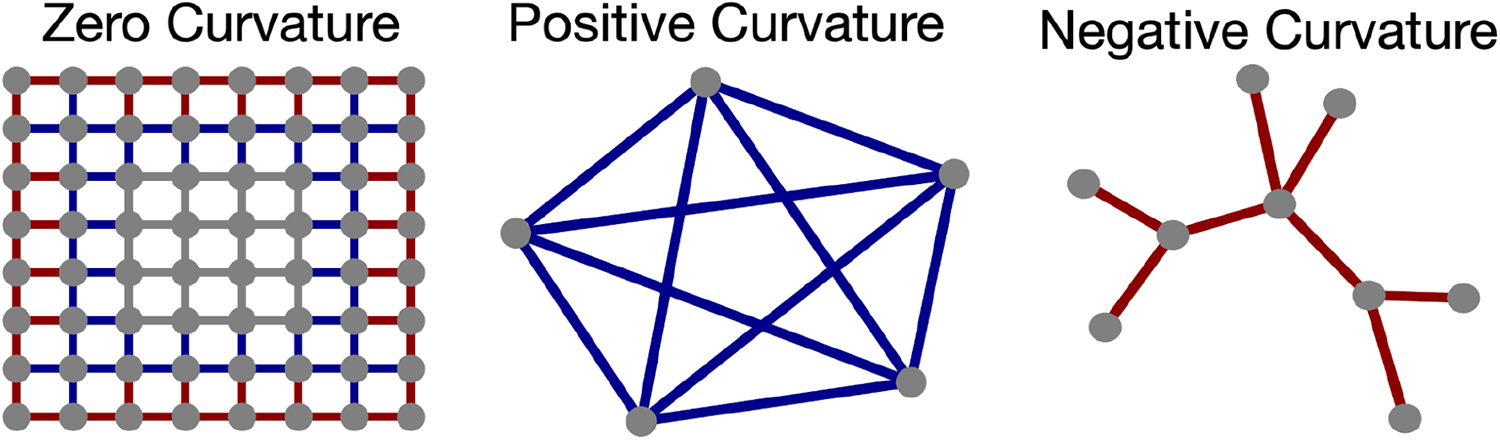
Ollivier Ricci curvature on example networks. Gray edges indicate zero curvature between nodes, blue edges indicate positive curvature, and red edges indicate negative curvature. In the center image, there are multiple paths that can be traced out between any pair of nodes; therefore, the curvature is positive. Conversely, the red edges in the right-most figure show negative curvature values since the removal of any edge would bisect the graph.

Formally, ORC is defined as follows:

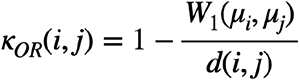

where W_i_ is the Wasserstein distance, also known as the Earth Mover’s distance (EMD), between the probability distributions, μ_i_, μ_j_. The probability distribution around a given node (gene), μ_i_, is defined by the edge weights originating from the given node *i* to adjacent nodes as follows:

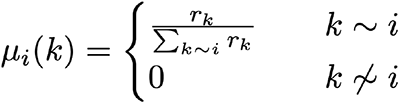

Where *r_k_* indicates either RNA-Seq or CNA values in node *k* connected to node *i.* The denominator *d(i, j)* is the weighted shortest path between the two nodes, where the edge weights of the weighted graph are derived from nodal values (RNA-Seq or CNA) quantifying the information between two nodes and is formally defined below.

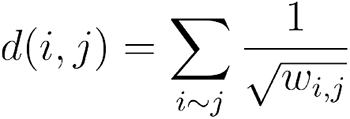

### Clustering analysis

To explore the potential subtypes in the cohort, we used a hierarchical agglomerative clustering method. For each data type, the RNA-Seq, CNA, and ORC matrices were separately clustered. The number of clusters was determined by the silhouette score (19), a measure which takes into account both the average intra-cluster distance and average nearest-cluster distance to determine the optimal number of clusters. Survival analysis for progression free survival (PFS) was performed using the Kaplan-Meier method and log-rank tests were used to determine statistical significance. Multiple comparisons were corrected using the Benjamini Hochberg false discovery rate (BH-FDR) (20).

### Differential gene expression analysis

To investigate biological differences between the identified subtypes, we conducted a differential gene expression analysis between high and low-risk groups, as identified in prior steps, using RNA sequencing read counts with DESeq2 (21). The p-values from this analysis were then BH-FDR corrected. Genes with a corrected p-value less than 0.05 and an absolute log2 fold change greater than 3.5 were considered significant.

### Pathway analysis

Pathway analysis was performed using the Broad Institute’s Gene Set Enrichment Analysis (GSEA) tool (22,23). The utilized pathways are from the hallmark gene set collection from the human molecular signatures database (MSigDB) (24). The fifty gene sets present different biological states and processes identified using manual curation. Gene association with the immune system was determined using ImmuneSigDB, an immune system pathways database provided by GSEA (25).

### Prognosis analysis

To test whether or not an individual gene was prognostic, we used a Cox’s proportional hazards model (26) with the RNA-Seq data. The p-values from this analysis were corrected for multiple hypothesis testing using BH-FDR. For genes that were significant with RNA-Seq, we repeated the modeling analysis using CNA data.

### Network topology analysis

To understand how genes are connected to each other, a given gene’s immediate neighbors are visualized as a ‘1-hop plot.’ Furthermore, a ‘2-hop plot’ shows not only a gene’s immediate neighbors but also the nearest neighbors of the immediate neighbor genes, in order to contextualize the relative portion of the overall network a given gene occupies. Bridge genes connect with relatively few genes in the network, while hub genes form many connections relative to the rest of the genes in the network.

### Data and code availability

The methods and instructions for how to use them are available for download at www.github.com/aksimhal/mm-orc-subtypes. All data is available for download at www.research.mmrf.org.

## RESULTS

### Patient cohort

CoMMpass IA19 RNA-Seq and CNA data were available for 659 patients. The mean age in the dataset was 62.5 ± 10.7 years; 60% were male, and the ISS distribution was 35% stage I, 35% stage II, and 30% stage III. For the cohort, the five-year PFS rate was approximately 32%, with the longest survival time listed at eight years. An overview is presented in **Supplementary Table 1**.

### Hierarchical clustering using Ollivier-Ricci curvature differentiates subtypes with low progression-free survival rates

The largest connected network component from shared information between the HPRD, RNA-Seq, and CNA data consisted of 8,468 nodes and 33,695 edges. ORC, a correlate for robustness of strength between gene interaction pairs, was computed for each of the 33,695 interaction pairs in each individual patient. Hierarchical clustering of the resultant ORC matrix together with CNA data produced 8 clusters (**Supplementary Figure 1A**, **Figure 3A**), while clustering based on RNA-Seq produced 6 clusters (**Supplementary Figure 1B**, **Figure 3B**); both methods being significant for PFS (CNA; p=0.0082, RNA-Seq; p=0.0016, log-rank test). Interestingly, the clustering appears to be defining biological differences not captured by the ISS prognostic score, with a relatively even distribution of ISS stages in each cluster.

Considering the dominant impact of hyperdiploidy on CNA analyses, we repeated hierarchical clustering on the non-hyperdiploid samples and found PFS prediction remained significant (p=0.0002, log-rank test). Of note, analyzing CNA via ORC produced a cluster representing 10% of patients with a markedly inferior PFS when compared to the remaining clusters (**Figures 3A, 3C**); median PFS was 1.7 years, despite only 35% of patients being ISS III. When assessing previously described copy number risk factors (**Supplementary Table 2**), patients in this cluster almost universally contain aberration in chr1q (gain; 57%, amplification; 29%, diploid 3%), while also harboring the highest proportion of the complex structural variant chromothripsis (43% of patients, p<0.0001 compared with the remaining clusters, Fisher’s exact test). This finding is congruent with previously published data demonstrating chromothripsis to be an independent prognostic factor in MM (17), and with an increasing body of knowledge demonstrating that multiple genomic insults compound to worse survival (17,27).

**Figure 3.**
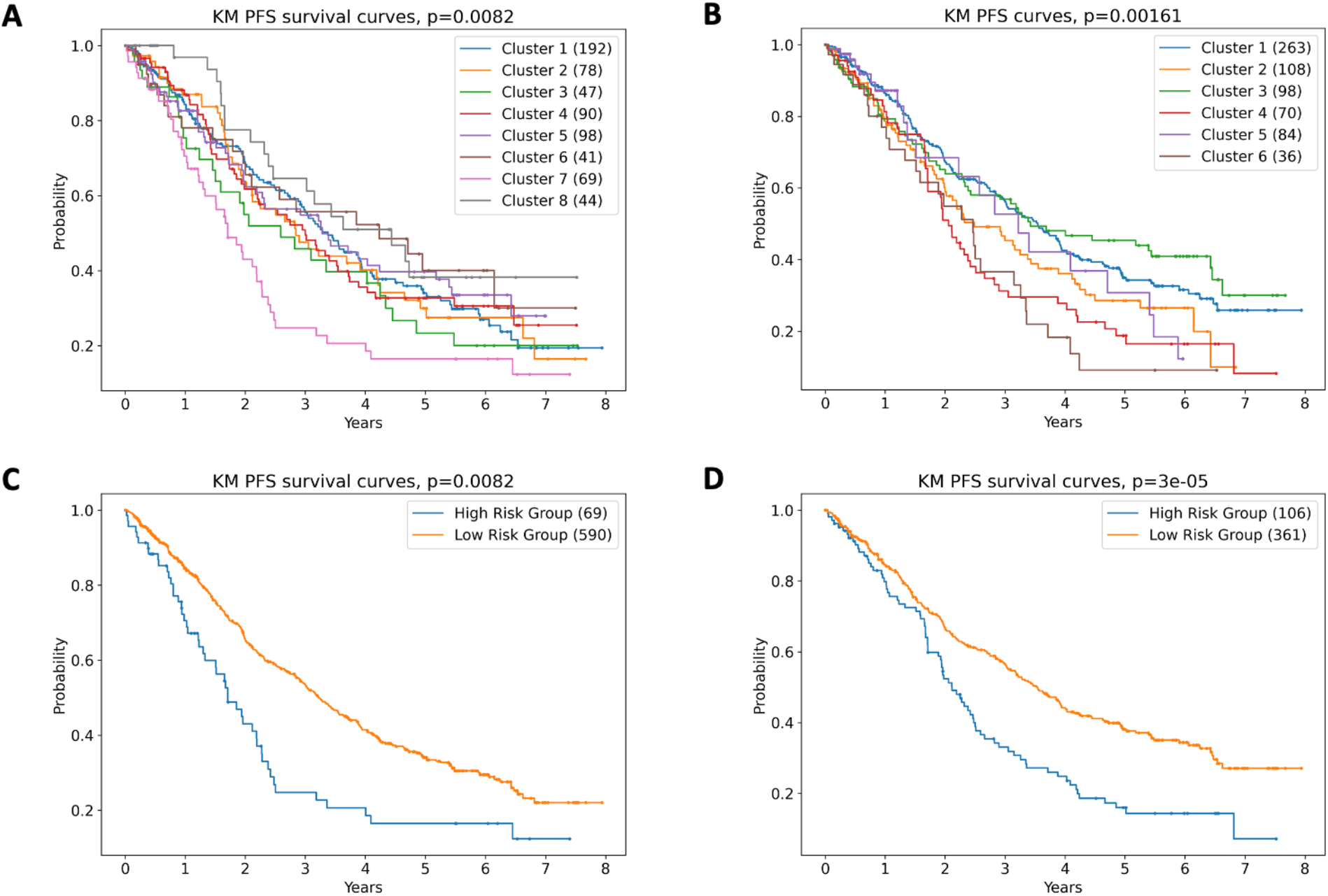
Hierarchical clustering using Ollivier Ricci Curvature (ORC) predicts progression-free survival (PFS) in multiple myeloma. Kaplan-Meier analysis of PFS based on ORC according to (A) copy number aberration, and (B) RNA sequencing. To better understand the differences between the high risk and low risk cohorts, clusters with similar outcomes were grouped. C) For CNA based clustering, clusters 1-6 and 8 were combined into the low-risk group. Cluster 7 was the high-risk group. D) For RNA-sequencing data, clusters 4 and 6 were combined into a high-risk group. Clusters 1 and 3 were combined into a low-risk group.

Clustering of the ORC matrix with RNA-Seq data produced more variation in PFS between clusters (**Figure 3B**). Of note, clusters 2 and 3 contain the majority of t(11;14) patients (**Supplementary Table 3**). Considering the dominant role of *CCND1* in MM pathophysiology, we repeated hierarchical clustering in the non-t(11;14) samples, which remained significant for PFS-prediction (p=0.0002, log-rank test). When clustering with all patients; 98% of those in cluster 4 harbor t(4;14), and 81% of those in cluster 6 have a translocation affecting *MAF,MAFA* or *MAFB*, with 72% having increased APOBEC-mutational activity. Clusters 1 and 5 are more heterogenous, with a combination of hyperdiploidy, canonical translocations, gain/amp1q, *TP53* aberration and chromothripsis. While a high proportion of patients in the 2 clusters with the shortest PFS (4 and 6) carry a previously described genomic risk factor, the other clusters (1 and 3) demonstrate a longer PFS despite 29.2% being ISS III, and 34% harboring a risk factor included in R-ISS / R2-ISS. Given that clustering with ORC using RNA-Seq demonstrated better discrimination of PFS compared with CNA, we have elected to focus on RNA-Seq for the remainder of the current study. We hypothesized that expanding on the ORC analysis with gene set enrichment analysis (GSEA), prognostic modeling, and network topology analysis will provide further biological insights.

### Expression analysis using ORC-based risk groups demonstrates differential DNA damage and immune system signaling

Differential gene expression analysis was conducted comparing high-risk (clusters 4 and 6) and low-risk (clusters 1 and 3) as defined by ORC analysis of RNA-Seq data. Gene sets enriched in the high-risk group includes inflammatory response, IL-6/JAK/STAT3 signaling and DNA damage response (DDR) signaling (P53 pathway, DNA repair and apoptosis, **Table 1**). Of note, there was no significant difference between the groups in p53 function by traditional methods (*TP53* mutations and del17p), therefore our methods are capturing more global dysregulation in DNA damage signaling than is evident by standard mutation and copy number analysis.

**Table 1.**
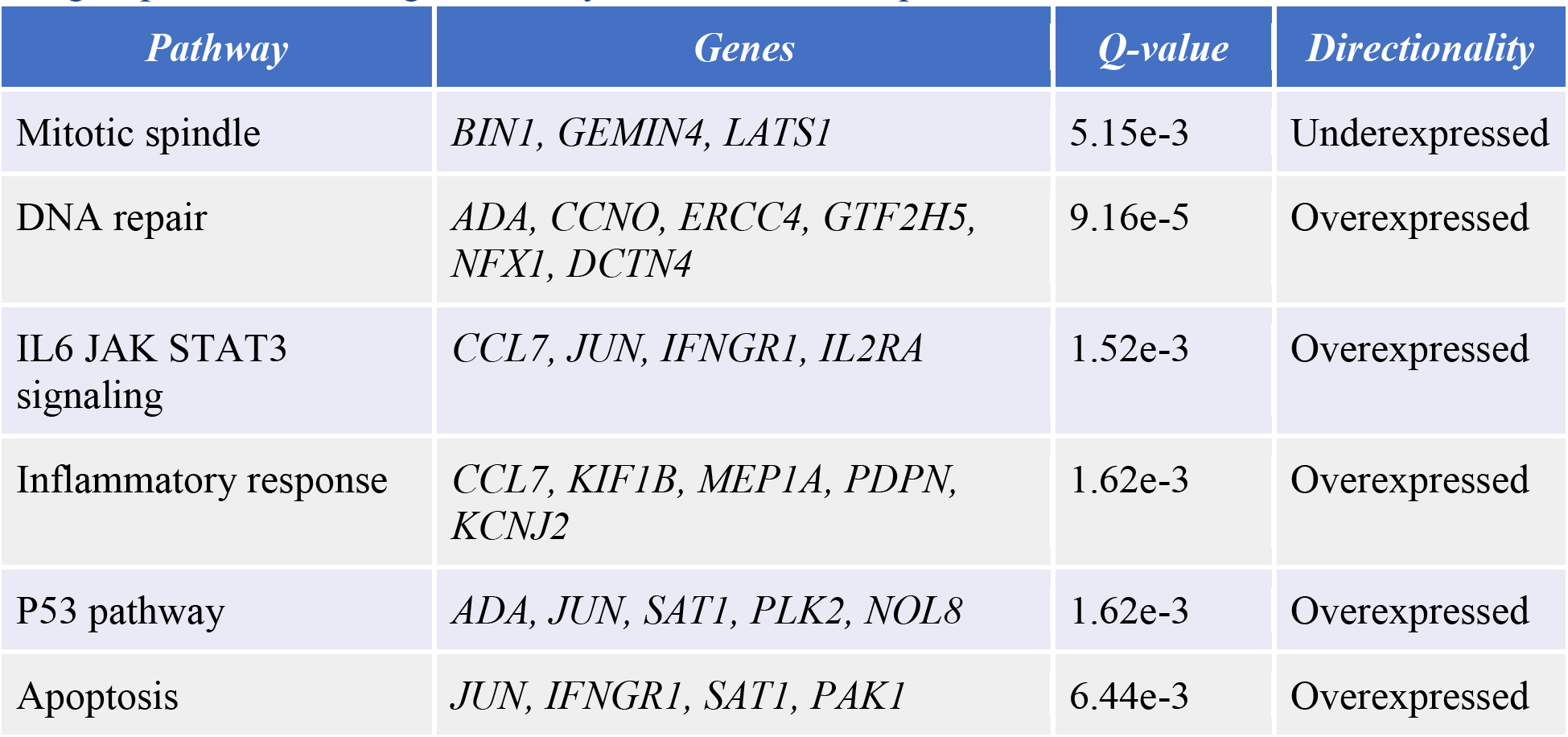
Differential gene expression analysis according to ORC-based risk groups. Directionality indicates the gene-set expression in the high-risk group compared with the low-risk group, with risk being defined by ORC of RNA-Seq data.

Within these differentially expressed pathways, 118 genes were selected for further pathway analysis (having absolute log fold change > 3.5 and corrected p-value < 0.05, **Supplementary Table 4**). Of these 118 genes, 19 were under-expressed and 99 were overexpressed in the short survival group compared to the longer survival group in the poor survival group. In univariate analysis, 8/118 genes were predictors of PFS (*BUB1, MCM1, NOSTRIN, PAM, RNF115, SNCAIP, SPRR2A* and *WEE1,* **Table 2**), with 5 of these also being significant when analyzing based on CNA (*NOSTRIN, PAM, RNF115, SNCAIP* and *SPRR2A*). Interestingly, none of these genes feature in previously described lists of MM driver genes (27,28), suggesting that we are capturing novel aspects of MM biology. In addition to differential expression in the inflammatory response and IL-6/JAK/STAT3 signaling gene sets, interrogation of the ImmuneSigDB database demonstrated 110 /118 genes to overlap with ImmuneSigDB pathways, including all 8 of the independently prognostic genes (**Table 2**). Taken together, these findings suggest that global assessment of gene interactions can detect complex immune dysregulation.

**Table 2.** Gene expression in 8 novel immune-network genes associate with survival. Coefficients less than 1 indicate a protective effect — associated with longer PFS. Coefficients greater than 1 indicate a detrimental effect — associated with a shorter PFS.

| *Gene* | *Coefficient* | *95% - 105% Range* | *Q-value* | *Gene description* | *Number of ImmunoSigDB gene sets* |
| --- | --- | --- | --- | --- | --- |
| *BUB1* | 1.36 ± 0.05 | 1.22-1.51 | 1.71e-8 | BUB1 mitotic checkpoint serine/threonine kinase | 5 |
| *MCM6* | 1.45 ± 0.07 | 1.27-1.66 | 6.19e-8 | Minichromosome maintenance complex component 6 | 4 |
| *NOSTRIN* | 1.58 ± 0.11 | 1.27-1.98 | 4.49e-5 | Nitric oxide synthase trafficking | 1 |
| *PAM* | 0.72 ± 0.08 | 0.62-0.83 | 1.34e-5 | Peptidylglycine alpha-amidating monooxygenase | 7 |
| *RNF115* | 1.42 ± 0.11 | 1.14-1.77 | 1.72e-3 | Ring finger protein 115 | 6 |
| *SNCAIP* | 1.40 ± 0.09 | 1.17-1.67 | 2.03e-4 | Synuclein alpha interacting protein | 1 |
| *SPRR2A* | 1.34 ± 0.05 | 1.22-1.46 | 1.43e-10 | Small proline rich protein 2A | 3 |
| *WEE1* | 1.32 ± 0.04 | 1.23-1.41 | 6.19e-15 | WEE1 G2 checkpoint kinase | 9 |

### Local neighborhood 1-hop and 2-hop gene networks demonstrate differential DNA damage and immune system signaling

A key feature of gene network analysis is the ability to capture a wide range of gene-pair interactions, above and beyond the expression levels of a single gene. While this analysis may be difficult to interpret in the context of highly connected genes, it can detect complex patterns (i.e., an overall increase or decrease in network robustness) or specific individual interactions (i.e., a gene-pair demonstrating an increase in robustness while all other local connections become more fragile).

Comparing high-risk and low-risk clusters as defined by ORC analysis of RNA-Seq data, we note several interesting network expression patterns. Within DDR-signaling, *TP53* and *ATM* signaling pathways overwhelming become more robust in the high-risk group (**Figures 4A, 4B**), with more robust pathways generally expected to exert increased effects. While we typically associate loss of p53 function with poor prognosis in cancer, global network analysis is detecting global changes in expression that may not fully capture functional protein levels. The same analysis performed on the basis of CNA demonstrates a mixture of *TP53* connections becoming more robust and more fragile, possibly reflecting the impact of del17p (**Supplementary Figure 2A**).

In addition to DDR-signaling, networks centered on *CCND1* and *MYC* become more robust overall (**Figures 4C, 4D**), which suggests these signaling and transcriptional hubs remain dominant in the context of high-risk disease. In contrast to the above networks showing a clear signal of robustness, the effect on *RAF / RAS / MAPK* and *NFKB* signaling are more heterogenous (**Supplementary Figure 2B-D**), suggesting that some parts of this network may play an oversized role in MM biology compared with the other interactions.

Considering the immune dysregulation observed on GSEA analysis, signaling through some cytokines and receptors become more fragile (i.e., IL-6, IFNg; **Figures 4E, 4F**), while others demonstrate a more heterogenous effect (i.e., TNF, IFNa; **Supplementary Figures 2F, 2G**). In this context, pathways becoming more fragile would be expected to exert less than normal control. Interestingly, multiple networks involving therapeutic targets for MM immune-based therapies become more fragile, suggesting potential therapeutic vulnerabilities. This included *TNFRSF17* (encoding for BCMA, a cellular-therapy target), *CD38* (the target of monoclonal antibody daratumumab), *IZKF3* (a target of immunomodulatory agent lenalidomide) and *SLAMF7* (the target of monoclonal antibody elotuzumab) (**Figures 4G-I, Supplementary Figures 2H, 2I**).

**Figure 4.**
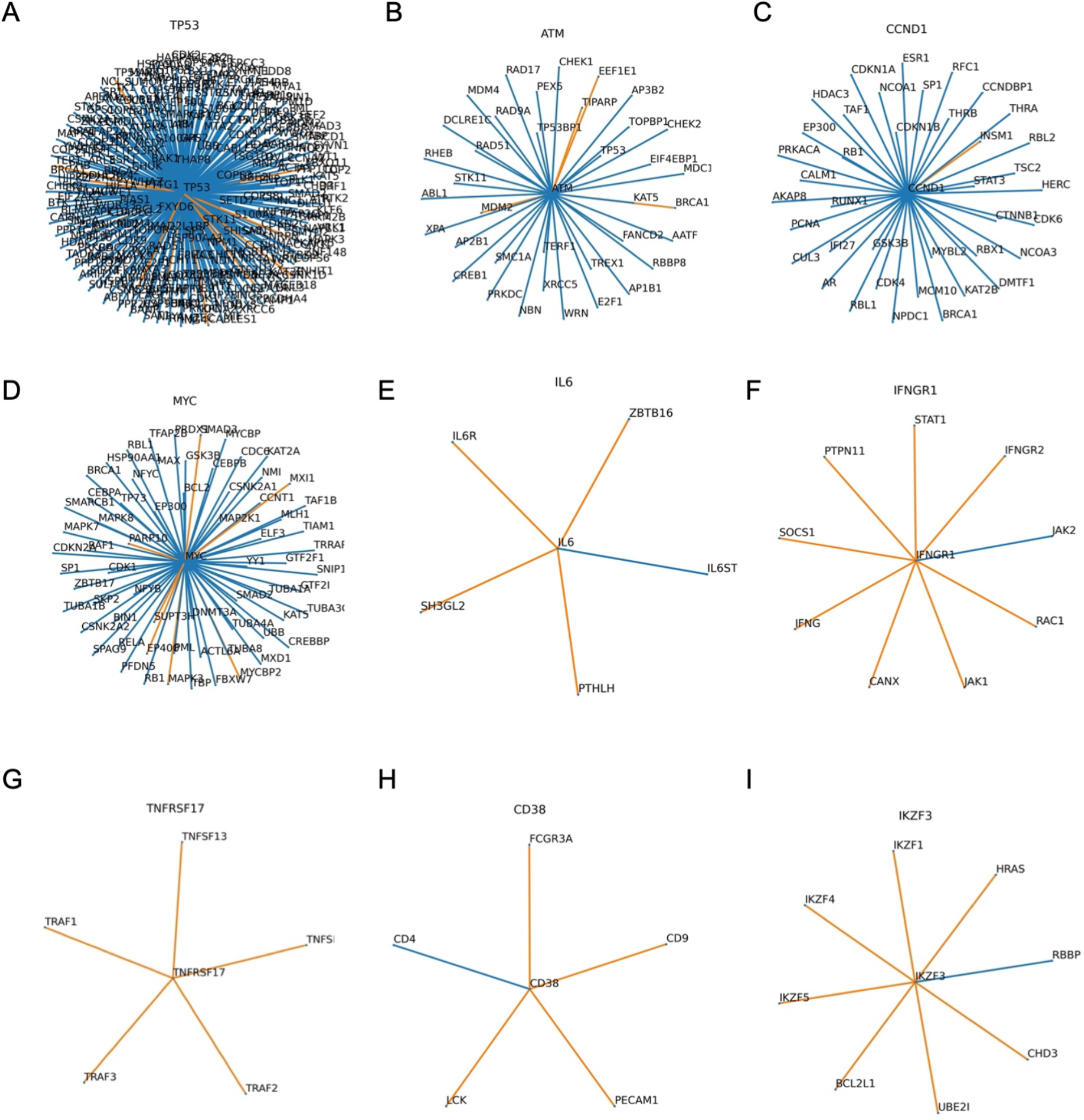
Local neighborhood of selected genes relevant to MM biology and the immune system. Each line or edge represents the interaction between a gene-pair in a network, comparing the median interactions observed in the high-risk group compared with those in the low-risk group. Blue edges indicate that the connections are more robust in the high-risk group, while orange edges are more fragile, risk being defined by the RNA-Seq-based clustering analysis. Higher resolution images are available at www.github.com/aksimhal/mm-orc-subtypes.

From the list of 8 novel genes having expression associated with PFS in MM, all have a recognized role in immune regulation (**Table 2**). In contrast with the other genes, only *WEE1, (*encoding for a tyrosine kinase which affects G2-M transition), has been previously implicated in MM biology (29). In the HPRD, *WEE1* acts as a hub gene, forming an above average number of connections with its immediate neighbors (18 versus 8.4 for the whole graph). Interestingly, within the 8 prognostic genes, *BUB1* and *WEE1* connect to each other in a 2-hop analysis via *PLK1*, *CDK1*, and *CRK*. From the genes with significantly different expression between risk groups, 24/118 (20.3%) connect to the 8 prognostic genes within the two-hop analysis.

The 8 genes identified play different roles in their local neighborhoods (**Figure 5**); *NOSTRIN*, (a nitric oxide synthase trafficker), *RNF115*, (an E3 ubiquitin ligase), and *SPRR2A* (induced by type-2 cytokines in response to infection) form bridge-like connections to a single other gene. *NOSTRIN* connects to another nitric oxide gene, *NOS3*, *RNF115* to the RAS oncogene family member *RAB7A*, while *SPRR2A* connects with *EVPL* (associated with squamous cell cancer and autoimmune disease). Four genes act as bridges for their local neighborhood: *BUB1*, *MCM6*, *PAM*, and *SNCAIP* (**Figures 5, 6**). While these genes are not hub genes per se, they connect to multiple hub genes and could therefore play a modulating role.

**Figure 5.**
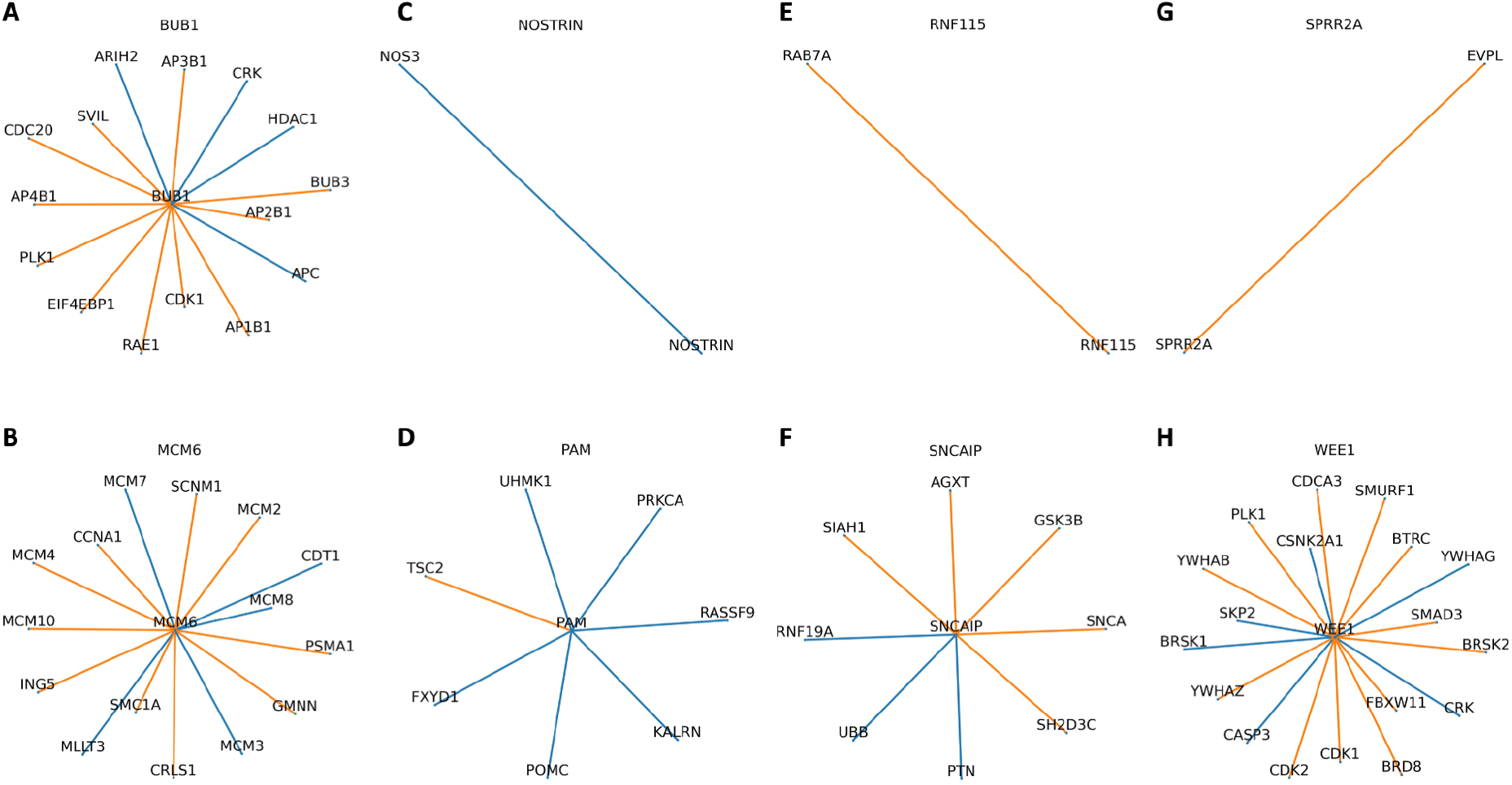
Local neighborhood of the eight genes identified as being predictive of PFS. Each line or edge represents the interaction between a gene-pair in a network, comparing the median interactions observed in the high-risk group compared with those in the low-risk group. Blue edges indicate that the connections are more robust in the high-risk group, while orange edges are more fragile, risk being defined by the RNA-Seq-based clustering analysis.

**Figure 6.**
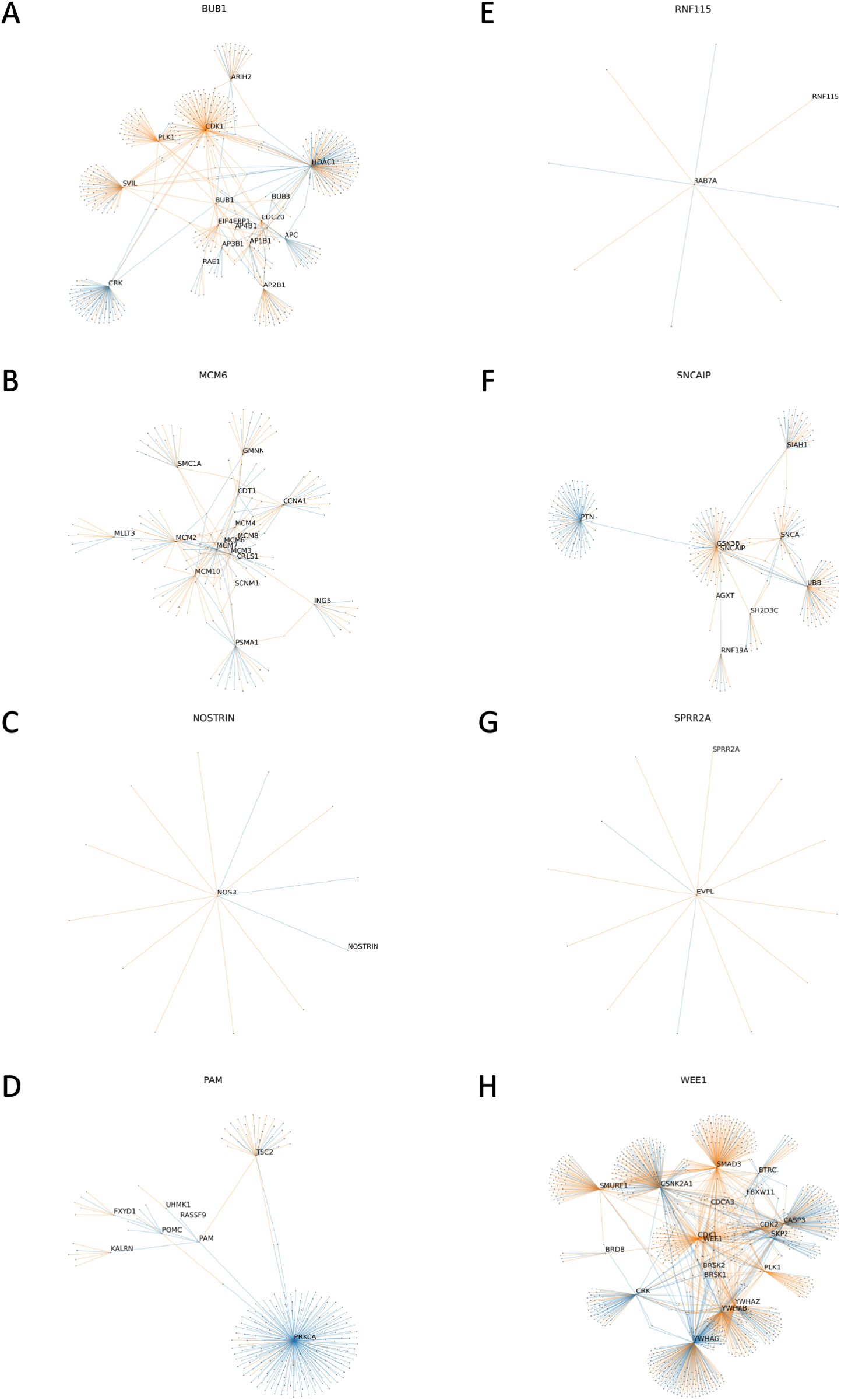
‘Two-hop’ neighborhood of the eight genes identified as being predictive of PFS. Each line or edge represents the interaction between a gene-pair in a network, comparing the median interactions observed in the high-risk group compared with those in the low-risk group. Blue edges indicate that the connections are more robust in the high-risk group, while orange edges are more fragile, risk being defined by the RNA-Seq-based clustering analysis. Higher resolution images are available at www.github.com/aksimhal/mm-orc-subtypes.

For example, in the 2-hop analysis, the mitotic checkpoint kinase *BUB1* connects to *HDAC1* (**Figure 6A**), a histone deacetylase commonly upregulated in MM cells with a well-defined impact on prognosis (30). We note multiple network connections between *BUB1* and *HDAC1*, as well as connections between *BUB1* and each of *CDK1* (cell-cycle transition regulator) and *APC* (a tumor-suppressor protein within the Wnt signaling pathway). *PAM*, encoding for a protein with multiple functions described, connects to *PRKCA,* a protein kinase involved in regulation of proliferation, tumorigenesis, and inflammation. Interestingly, the network connections around *PRKCA* are predominantly more robust in the high-risk group. *SNCAIP*, (which inhibits ubiquitin ligase activity), connects with *PTN (***Figure 6F**; a hub gene encoding for a protein having a role in cell survival, angiogenesis and tumorigenesis), previously noted to be elevated in MM patients (31). Our analysis finds that the connection between *SNCAIP* and *PTN* becomes more robust in the high-risk group. Interestingly, when comparing the 1- and 2-hop networks between RNA-Seq and CNA data, several gene networks were highly analogous between the two methods (Supplementary Figure 3).

Overall, the complex gene interactions captured through ORC analysis have the capacity to significantly improve our understanding of biological differences between patients have short and long survival, extending on what we understand from traditional mutation and copy number analysis.

## DISCUSSION

In order to investigate global gene-protein interaction networks in MM and their impact on prognosis, we combined a known protein interaction network, HPRD, with a large MM dataset; CoMMpass. We applied a novel measure of network robustness, ORC, to examine patterns in the RNA-Seq gene expression and CNA data and how they relate to clinical outcomes. Hierarchical clustering using ORC produced 6 clusters based on RNA-Seq and 8 clusters based on CNA data, with both data sources predictive of PFS. Previously published genomic classifications in MM based on RNA-Seq and/or CNA data have defined between four to twelve clusters, depending on the data and analytical approach (5–10). To date, no study has integrated genomic information with known protein interaction information in an analysis able to simultaneously integrate local and global network information. By using techniques previously shown to uncover differences in network strength in other domains, such as ovarian cancer and autism spectrum disorders (12,13), we were able to demonstrate a new way of characterizing MM genomic data.

Our results demonstrate fidelity with known genomic risk factors (i.e., t(4;14), gain 1q, *TP53* aberration) as well as emerging factors not yet in clinical use (i.e., APOBEC mutational activity and the complex structural variant chromothripsis (17,32,33). While some genomic subgroups were defined by a single event (i.e., 98% of RNA-Seq cluster 4 harboring t(4;14), the network analysis approach produced other groups not previously described, with a combination of genomic events defining prognostically significant clusters. It is notable that the cluster having the shortest PFS was defined not by ISS, R-ISS, hyperdiploidy or IgH translocations but associated with the combination of gain/amp 1q and chromothripsis. This finding supports the hypothesis that more comprehensive, global genomic characterization is able to better define MM prognosis.

As ORC measures relative robustness between genes, GSEA analysis comparing high-risk and low-risk groups as identified by ORC analysis of RNA-Seq data allowed exploration of gene-pair interaction changes in robustness associated with survival differences between groups. GSEA located 118 differentially expressed genes associated with six key biological pathways, five of which were overexpressed in the group with the poor survival. The underexpressed pathway, mitotic spindle assembly, has previously been reported to be associated with poor prognosis in MM (34), while the overexpressed pathways were all associated with DNA damage response (DDR) and acute phase inflammation / immune response. While del 17p is included in the R-ISS prognostic score, and genomic complexity and instability are recognized features of high-risk MM biology (35–38), there is not currently any immune component to routine prognostication of NDMM patients. Furthermore, there is likely a biological link between the pathways we describe, with an inflammatory hypoxic microenvironment potentially contributing to aberrant DDR (39), and functional high-risk patients who relapse within 12 months described to harbor both mutations affecting the IL-6/JAK/STAT pathway and abnormal gene expression associated with mitosis / DDR (40).

Univariate analysis of the 118 differentially expressed genes identified 8 prognostic genes which are all associated with immune function according to ImmunoSigDB. Network topology analysis identified most of these 8 to be bridge genes, connecting to genes known to have biological impact in MM (i.e., *HDAC1*, *CDK1*, *PRKCA* and *PTN*). The near-neighbor and 2-hop gene topology networks capture more global gene dysregulation, potentially missed in single-gene expression analysis. Our results may also suggest a new set of therapeutic targets to further investigate in high-risk MM patients.

Considering possible limitations; while CoMMpass represents the largest multi-site, international genomic MM dataset compiled to date, it does contain patients who received drug regimens no longer in common usage, and a low proportion of patients receiving the most potent modern regimens. Ideally our methods would be applied to datasets including daratumumab-based induction therapy. Considering possible extension of our analytical methods: while the choice of using the HPRD as the protein interaction network is common in literature (41), other networks, such as STRING (42), may provide complementary results. Finally, no network analysis method represents the ‘gold standard’, and it is plausible that other clustering and network analysis methods may provide alternative results. Future studies may consider whether or not the 118 genes associated with high-risk individuals are dysregulated at precursor MM stages, and how the expression of these genes is altered in response to treatment.

## Acknowledgments

The authors would like to thank Jonathan J. Keats at the Translational Genomics Research Institute and the team at the Multiple Myeloma Research Foundation for their incredible work and support with the CoMMpass dataset. KHM received support from the Multiple Myeloma Research Foundation, the American Society of Hematology, and the Royal Australasian College of Physicians. AKS, RE, JOD, JHO, and AT are supported by the Breast Cancer Research Foundation. SZU is supported by both the Leukemia Lymphoma Society and International Myeloma Society.

## Author contributions

AKS: conceptualization, formal analysis, data curation, visualization, writing-original draft. KHM: conceptualization, supervision, writing-original draft, writing-review and editing. RE: methodology, writing – original draft. JZ: methodology, writing – original draft. SZU: conceptualization, writing-review and editing. LN: conceptualization, supervision, writing-review and editing. JOD: conceptualization, supervision, writing-review and editing. JHO: conceptualization, supervision, writing-review and editing. AT: conceptualization, supervision, writing-review and editing.

## Data availability

The datasets used are available for download at http://research.themmrf.org.

## References

1. Morgan GJ, Walker BA, Davies FE. The genetic architecture of multiple myeloma. Nat Rev Cancer. 2012 Apr 12;12(5):335–48.

2. Hu Y, Chen W, Wang J. Progress in the identification of gene mutations involved in multiple myeloma. Onco Targets Ther. 2019 May 24;12:4075–80.

3. Palumbo A, Avet-Loiseau H, Oliva S, Lokhorst HM, Goldschmidt H, Rosinol L, et al. Revised International Staging System for multiple myeloma: A report from international myeloma working group. J Clin Oncol. 2015 Sep 10;33(26):2863–9.

4. D’Agostino M, Cairns DA, Lahuerta JJ, Wester R, Bertsch U, Waage A, et al. Second Revision of the International Staging System (R2-ISS) for overall survival in multiple myeloma: A European Myeloma Network (EMN) report within the HARMONY project. J Clin Oncol. 2022 Oct 10;40(29):3406–18.

5. Zhan F, Huang Y, Colla S, Stewart JP, Hanamura I, Gupta S, et al. The molecular classification of multiple myeloma. Blood. 2006 Sep 15;108(6):2020–8.

6. Chng WJ, Kumar S, Vanwier S, Ahmann G, Price-Troska T, Henderson K, et al. Molecular dissection of hyperdiploid multiple myeloma by gene expression profiling. Cancer Res. 2007 Apr 1;67(7):2982–9.

7. Broyl A, Hose D, Lokhorst H, de Knegt Y, Peeters J, Jauch A, et al. Gene expression profiling for molecular classification of multiple myeloma in newly diagnosed patients. Blood. 2010 Oct 7;116(14):2543–53.

8. Jang JS, Li Y, Mitra AK, Bi L, Abyzov A, van Wijnen AJ, et al. Molecular signatures of multiple myeloma progression through single cell RNA-Seq. Blood Cancer J. 2019 Jan 3;9(1):2.

9. 9. Skerget S, Penaherrera D, Chari A, Jagannath S, Siegel DS, Vij R, et al. Genomic Basis of Multiple Myeloma Subtypes from the MMRF CoMMpass Study [Internet]. bioRxiv. medRxiv; 2021. Available from: http://medrxiv.org/lookup/doi/10.1101/2021.08.02.21261211

10. Bustoros M, Anand S, Sklavenitis-Pistofidis R, Redd R, Boyle EM, Zhitomirsky B, et al. Genetic subtypes of smoldering multiple myeloma are associated with distinct pathogenic phenotypes and clinical outcomes. Nat Commun. 2022 Jun 15;13(1):3449.

11. Sandhu RS, Georgiou TT, Reznik E, Zhu L, Kolesov I, Senbabaoglu Y, et al. Graph curvature for differentiating cancer networks. Sci Rep. 2015;5:12323.

12. Elkin R, Oh JH, Liu YL, Selenica P, Weigelt B, Reis-Filho JS, et al. Geometric network analysis provides prognostic information in patients with high grade serous carcinoma of the ovary treated with immune checkpoint inhibitors. NPJ Genom Med. 2021 Nov 24;6(1):99.

13. Simhal AK, Carpenter KLH, Kurtzberg J, Song A, Tannenbaum A, Zhang L, et al. Changes in the geometry and robustness of diffusion tensor imaging networks: Secondary analysis from a randomized controlled trial of young autistic children receiving an umbilical cord blood infusion. Front Psychiatry [Internet]. 2022;13. Available from: https://www.frontiersin.org/articles/10.3389/fpsyt.2022.1026279

14. Keats JJ, Craig DW, Liang W, Venkata Y, Kurdoglu A, Aldrich J, et al. Interim analysis of the mmrf CoMMpass trial, a longitudinal study in multiple myeloma relating clinical outcomes to genomic and immunophenotypic profiles. Blood. 2013 Nov 15;122(21):532– 532.

15. Peri S, Navarro JD, Kristiansen TZ, Amanchy R, Surendranath V, Muthusamy B, et al. Human protein reference database as a discovery resource for proteomics. Nucleic Acids Res. 2004 Jan 1;32(Database issue):D497–501.

16. Patro R, Duggal G, Love MI, Irizarry RA, Kingsford C. Salmon provides fast and bias-aware quantification of transcript expression. Nat Methods. 2017 Apr;14(4):417–9.

17. Rustad EH, Yellapantula VD, Glodzik D, Maclachlan KH, Diamond B, Boyle EM, et al. Revealing the impact of structural variants in multiple myeloma. Blood Cancer Discov. 2020 Nov;1(3):258–73.

18. Ollivier Y. Ricci curvature of metric spaces. C R Math. 2007;345(11):643–6.

19. Rousseeuw PJ. Silhouettes: A graphical aid to the interpretation and validation of cluster analysis. J Comput Appl Math. 1987 Nov 1;20:53–65.

20. Benjamini Y, Hochberg Y. Controlling the false discovery rate: a practical and powerful approach to multiple testing. J R Stat Soc [Internet]. 1995; Available from: https://rss.onlinelibrary.wiley.com/doi/abs/10.1111/j.2517-6161.1995.tb02031.x

21. Love MI, Huber W, Anders S. Moderated estimation of fold change and dispersion for RNA-seq data with DESeq2. Genome Biol. 2014;15(12):550.

22. Subramanian A, Tamayo P, Mootha VK, Mukherjee S, Ebert BL, Gillette MA, et al. Gene set enrichment analysis: a knowledge-based approach for interpreting genome-wide expression profiles. Proc Natl Acad Sci U S A. 2005 Oct 25;102(43):15545–50.

23. Mootha VK, Lindgren CM, Eriksson K-F, Subramanian A, Sihag S, Lehar J, et al. PGC-1alpha-responsive genes involved in oxidative phosphorylation are coordinately downregulated in human diabetes. Nat Genet. 2003 Jul;34(3):267–73.

24. Liberzon A, Birger C, Thorvaldsdóttir H, Ghandi M, Mesirov JP, Tamayo P. The Molecular Signatures Database (MSigDB) hallmark gene set collection. Cell Syst. 2015 Dec 23;1(6):417–25.

25. Godec J, Tan Y, Liberzon A, Tamayo P, Bhattacharya S, Butte AJ, et al. Compendium of immune signatures identifies conserved and species-specific biology in response to inflammation. Immunity. 2016 Jan 19;44(1):194–206.

26. Royston P, Parmar MKB. Flexible parametric proportional-hazards and proportional-odds models for censored survival data, with application to prognostic modelling and estimation of treatment effects. Stat Med. 2002 Aug 15;21(15):2175–97.

27. Walker BA, Mavrommatis K, Wardell CP, Ashby TC, Bauer M, Davies F, et al. A high-risk, Double-Hit, group of newly diagnosed myeloma identified by genomic analysis. Leukemia. 2019 Jan;33(1):159–70.

28. Maura F, Bolli N, Angelopoulos N, Dawson KJ, Leongamornlert D, Martincorena I, et al. Genomic landscape and chronological reconstruction of driver events in multiple myeloma. Nat Commun. 2019 Aug 23;10(1):3835.

29. Barbosa RSS, Dantonio PM, Guimarães T, de Oliveira MB, Fook Alves VL, Sandes AF, et al. Sequential combination of bortezomib and WEE1 inhibitor, MK-1775, induced apoptosis in multiple myeloma cell lines. Biochem Biophys Res Commun. 2019 Nov 12;519(3):597–604.

30. Mithraprabhu S, Kalff A, Chow A, Khong T, Spencer A. Dysregulated Class I histone deacetylases are indicators of poor prognosis in multiple myeloma. Epigenetics. 2014 Nov;9(11):1511–20.

31. Yeh HS, Chen H, Manyak SJ, Swift RA, Campbell RA, Wang C, et al. Serum pleiotrophin levels are elevated in multiple myeloma patients and correlate with disease status. Br J Haematol. 2006 Jun;133(5):526–9.

32. Walker BA, Wardell CP, Murison A, Boyle EM, Begum DB, Dahir NM, et al. APOBEC family mutational signatures are associated with poor prognosis translocations in multiple myeloma. Nat Commun. 2015 Apr 23;6:6997.

33. Maura F, Petljak M, Lionetti M, Cifola I, Liang W, Pinatel E, et al. Biological and prognostic impact of APOBEC-induced mutations in the spectrum of plasma cell dyscrasias and multiple myeloma cell lines. Leukemia. 2018 Apr;32(4):1044–8.

34. Tao Y, Yang G, Yang H, Song D, Hu L, Xie B, et al. TRIP13 impairs mitotic checkpoint surveillance and is associated with poor prognosis in multiple myeloma. Oncotarget. 2017 Apr 18;8(16):26718–31.

35. Kassambara A, Gourzones-Dmitriev C, Sahota S, Rème T, Moreaux J, Goldschmidt H, et al. A DNA repair pathway score predicts survival in human multiple myeloma: the potential for therapeutic strategy. Oncotarget. 2014 May 15;5(9):2487–98.

36. Ali JYH, Fitieh AM, Ismail IH. The Role of DNA Repair in Genomic Instability of Multiple Myeloma. Int J Mol Sci [Internet]. 2022 May 19;23(10). Available from: http://dx.doi.org/10.3390/ijms23105688

37. Giesen N, Paramasivam N, Toprak UH, Huebschmann D, Xu J, Uhrig S, et al. Comprehensive genomic analysis of refractory multiple myeloma reveals a complex mutational landscape associated with drug resistance and novel therapeutic vulnerabilities. Haematologica. 2022 Aug 1;107(8):1891–901.

38. Maura F, Boyle EM, Rustad EH, Ashby C, Kaminetzky D, Bruno B, et al. Chromothripsis as a pathogenic driver of multiple myeloma. Semin Cell Dev Biol. 2022 Mar;123:115–23.

39. Saitoh T, Oda T. DNA damage response in multiple myeloma: The role of the tumor microenvironment. Cancers (Basel). 2021 Jan 28;13(3):504.

40. Soekojo CY, Chung T-H, Furqan MS, Chng WJ. Genomic characterization of functional high-risk multiple myeloma patients. Blood Cancer J. 2022 Jan 31;12(1):24.

41. Wu J, Vallenius T, Ovaska K, Westermarck J, Mäkelä TP, Hautaniemi S. Integrated network analysis platform for protein-protein interactions. Nat Methods. 2009 Jan;6(1):75– 7.

42. von Mering C, Huynen M, Jaeggi D, Schmidt S, Bork P, Snel B. STRING: a database of predicted functional associations between proteins. Nucleic Acids Res. 2003 Jan 1;31(1):258–61.

