## Supplementary Information for "Gene interaction network analysis in multiple myeloma detects complex immune dysregulation associated with shorter survival"

**Supplementary Figure 1. Average silhouette score according to number of clusters.**

The average silhouette score for hierarchical clustering based on (A) copy number aberration and (B) RNA-sequencing defined the optimal number of clusters to be 8 for copy number and 6 for RNA-sequencing.

A

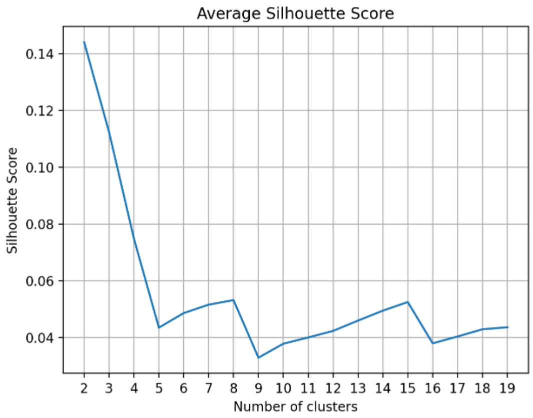


B

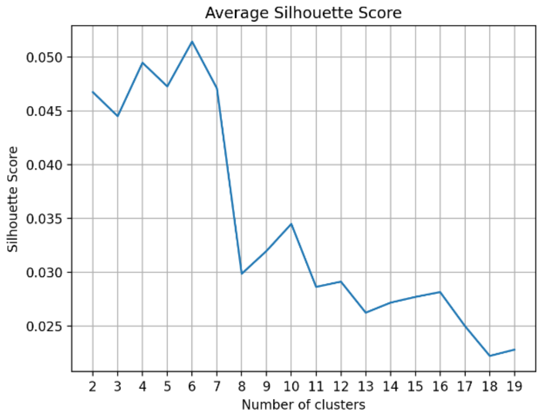

**Supplementary Figure 2. Near-neighbor analysis of gene-interactions for selected genes of significance in multiple myeloma biology based on copy number aberration.**

Each line or edge represents the interaction between a gene-pair in a network, comparing the median interactions observed in the high-risk group compared with those in the low-risk group. Blue edges indicate that the connections are more robust in the high-risk group, while orange edges are more fragile, risk being defined by the RNA-seq-based clustering analysis.

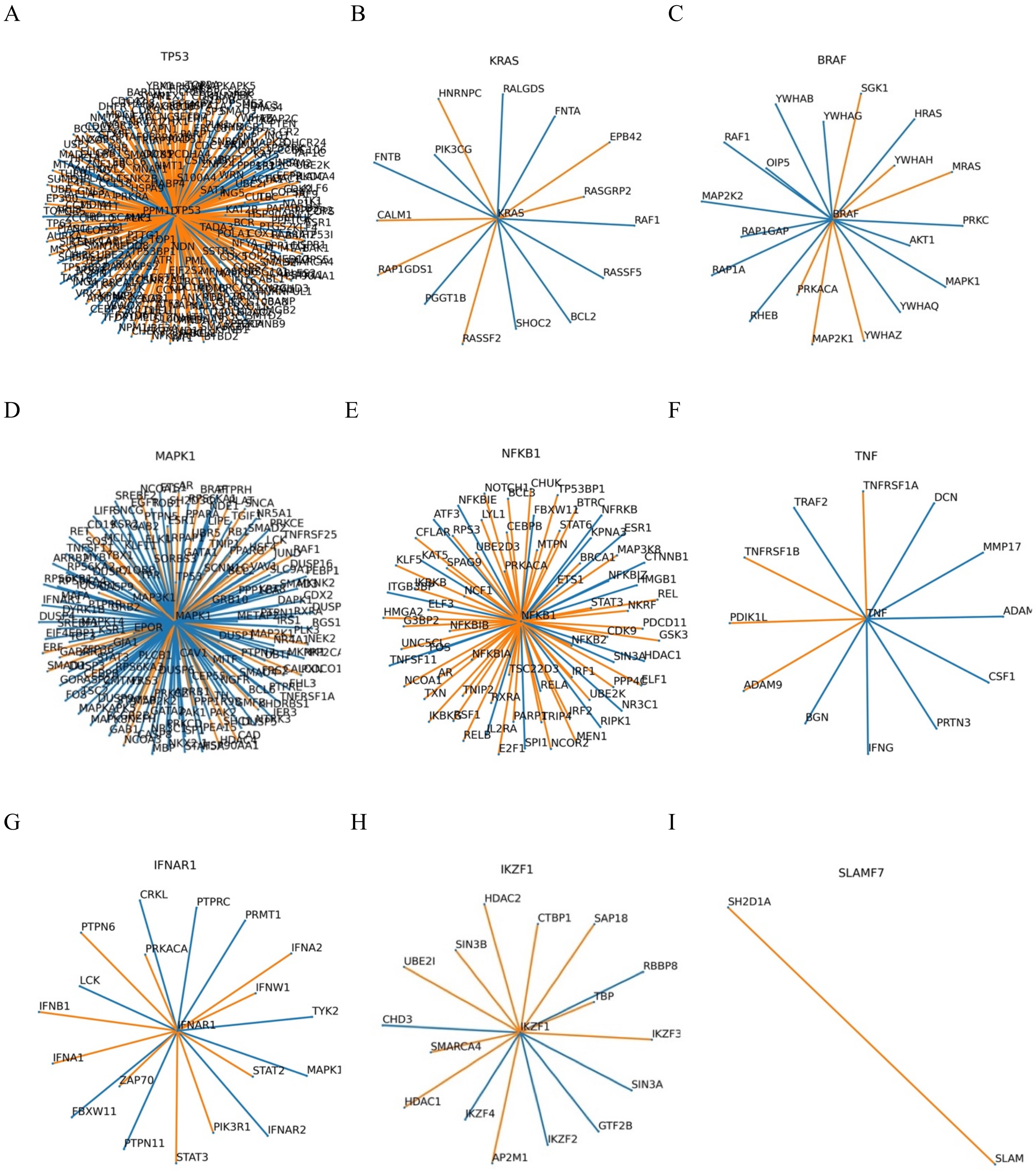

**Supplementary Figure 3. Near-neighbor analysis of gene-interactions for genes of prognostic significance in multiple myeloma biology based on copy number aberration.**

*PAM* presented as (A) 1-hop and (B) 2-hop network, *RNF115* presented as (C) 1-hop and (D) 2-hop network, and (E) *SNCAIP* 2-hop network. Each line or edge represents the interaction between a gene-pair in a network, comparing the median interactions observed in the high-risk group compared with those in the low-risk group. Blue edges indicate that the connections are more robust in the high-risk group, while orange edges are more fragile, risk being defined by the RNA-seq-based clustering analysis.

A

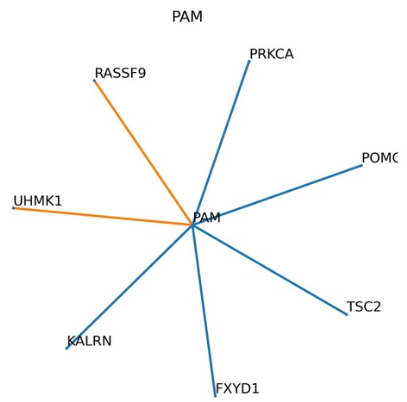


B

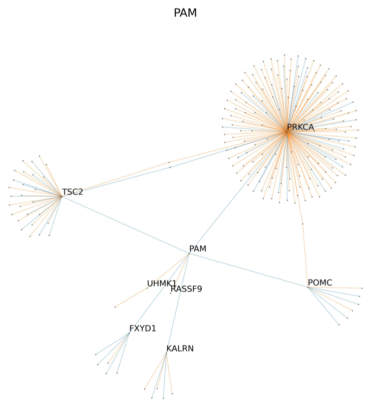


C

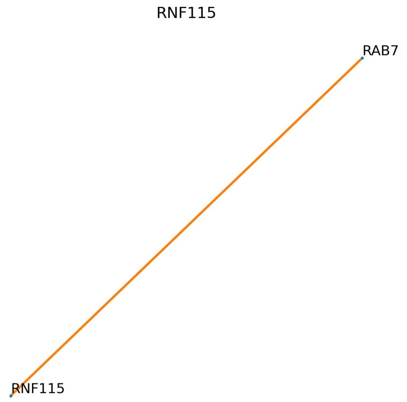


D

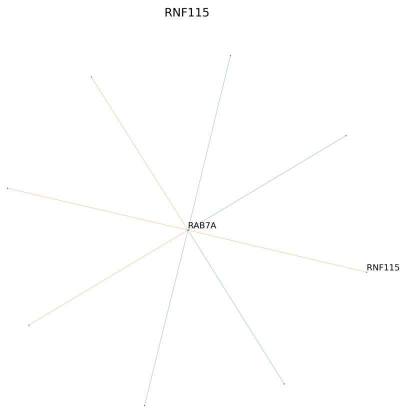


E

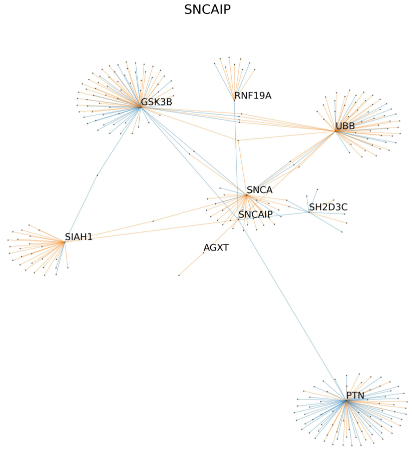

**Supplementary Table 1. Patient characteristics.**

| Variables | Patients (N=659) |
|---|---|
| Sex | Male: 393<br>Female: 266 |
| Age (mean ± SD) | 61.58 ± 13.09 years |
| ISS | Stage I: 229<br>Stage II: 232<br>Stage III: 179 |
| Treatment class | Bortezomib-based: 138<br>Carfilzomib-based: 38<br>IMIDs-based: 32<br>Combined bortezomib/IMIDs-based: 317<br>Combined IMIDs/carfilzomib-based: 110<br>Other: 24 |

**Supplementary Table 2. Patient clusters based on copy number aberration.**

| Patient cluster | 1 | 2 | 3 | 4 | 5 | 6 | 7 | 8 |
|---|---|---|---|---|---|---|---|---|
| **Hyperdiploidy** | 145/152 | 13/65 | 2/36 | 83/85 | 3/80 | 34/36 | 7/52 | 26/37 |
| **t(4:14)** | 3/168 | 24/70 | 11/37 | 2/88 | 6/82 | 1/38 | 24/58 | 8/38 |
| **t(11;14)** | 6/168 | 5/70 | 17/37 | 2/88 | 65/82 | 1/38 | 13/58 | 1/38 |
| **Translocation involving *MAF/MAFA/MAFB*** | 3/168 | 2/70 | 6/37 | 1/88 | 4/82 | 1/38 | 15/58 | 2/38 |
| **Translocation involving *MYC*** | 32/168 | 11/70 | 5/37 | 24/88 | 3/82 | 6/38 | 11/58 | 4/38 |
| **Chromothripsis** | 48/168 | 11/70 | 8/37 | 21/88 | 12/82 | 7/38 | 25/58 | 15/38 |
| **gain1q21** | (0): 139; (1): 12; (2): 1 | (0): 41; (1): 21; (2): 3 | (0): 26; (1): 5; (2): 5 | (0): 43; (1): 40; (2): 2 | (0): 78; (1): 2 | (0): 24; (1): 12 | (0): 2; (1): 33; (2): 17 | (0): 18; (1): 17; (2): 2 |
| ***TP53* aberration** | (0): 113; (1): 16; (2): 7 | (0): 54; (1): 5 | (0): 28; (1): 1; (2): 3 | (0): 71; (1): 11 | (0): 56; (1): 7; (2): 5 | (0): 29 | (0): 43; (1): 5; (2): 1 | (0): 24; (1): 8; (2): 2 |
| **Hyper-APOBEC** | 6/167 | 3/70 | 8/36 | 1/88 | 5/81 | 3/38 | 11/58 | 3/38 |

**Key:** for gain 1q21, 0 = diploid, 1 = gain (3 copies), 2 = amplification (4 or more copies). For *TP53* aberration, 0 = diploid, 1 = either deletion or mutation, 2 = biallelic loss.

**Supplementary Table 3. Patient clusters based on RNA sequencing.**

| Patient cluster | 1 | 2 | 3 | 4 | 5 | 6 |
|---|---|---|---|---|---|---|
| **Hyperdiploidy** | 197/229 | 62/93 | 9/82 | 12/63 | 28/46 | 5/30 |
| **t(4:14)** | 0/247 | 2/98 | 0/85 | 64/65 | 11/52 | 2/32 |
| **t(11;14)** | 3/247 | 19/98 | 77/85 | 0/65 | 11/52 | 0/32 |
| **Translocation involving *MAF/MAFA/MAFB*** | 1/247 | 2/98 | 0/85 | 2/65 | 3/52 | 26/32 |
| **Translocation involving *MYC*** | 61/247 | 13/98 | 4/85 | 7/65 | 7/52 | 4/32 |
| **Chromothripsis** | 62/247 | 21/98 | 7/85 | 26/65 | 17/52 | 14/32 |
| **gain1q21** | (0): 154; (1): 66; (2): 9 | (0): 70; (1): 21; (2): 2 | (0): 72; (1): 10 | (0): 33; (1): 18; (2): 12 | (0): 30; (1): 15; (2): 1 | (0): 12; (1): 12; (2): 6 |
| ***TP53* aberration** | (0): 178; (1): 24; (2): 5 | (0): 69; (1): 9; (2): 3 | (0): 60; (1): 5; (2): 4 | (0): 49; (1): 7; (2): 3 | (0): 40; (1): 4; (2): 1 | (0): 22; (1): 4; (2): 2 |
| **Hyper-APOBEC** | 4/247 | 7/97 | 2/83 | 1/65 | 3/52 | 23/32 |

**Key:** for gain 1q21, 0 = diploid, 1 = gain (3 copies), 2 = amplification (4 or more copies). For *TP53* aberration, 0 = diploid, 1 = either deletion or mutation, 2 = biallelic loss.

**Supplementary Table 4. 118 genes with significantly differential expression between high-risk and low-risk patient clusters.**

| Gene | FDR-BH corrected p value | log2FoldChange |
|---|---|---|
| ACTB | 8.15E-09 | -3.9791242 |
| ADA | 6.49E-49 | -3.6033571 |
| ANAPC1 | 2.29E-51 | -3.6378753 |
| ANKS3 | 2.18E-34 | -3.8578852 |
| APBB2 | 7.39E-60 | 4.17615932 |
| ATP6AP1 | 4.47E-14 | -4.0398878 |
| ATXN7L3 | 4.02E-58 | -5.0704603 |
| BACH2 | 0 | -11.442387 |
| BCL9 | 2.20E-37 | -4.7917991 |
| BIN1 | 1.61E-12 | 3.95216124 |
| BRSK2 | 2.74E-104 | -6.6614015 |
| BUB1 | 3.01E-42 | -3.7545566 |
| C16orf74 | 3.61E-25 | -3.5439076 |
| C20orf27 | 5.64E-46 | -4.2841877 |
| CACNA1D | 8.33E-53 | -4.8883489 |
| CAPNS1 | 3.16E-23 | -4.3470666 |
| CBX4 | 8.72E-23 | -3.9309193 |
| CCL7 | 1.32E-70 | -4.7256429 |
| CCNO | 3.78E-30 | -4.9069896 |
| CD4 | 7.77E-75 | 4.71988701 |
| CDK18 | 9.11E-66 | -3.8441406 |
| CEACAM5 | 2.02E-31 | -3.6537205 |
| CHD3 | 1.80E-51 | -3.5925837 |
| CMPK1 | 3.26E-36 | -4.6679916 |
| COL4A3 | 1.06E-35 | -3.6036391 |
| COL4A5 | 4.72E-51 | 3.84907315 |
| COPS6 | 4.68E-163 | -6.6641661 |
| CUEDC1 | 1.21E-10 | -4.0609393 |
| CYP17A1 | 2.31E-67 | -3.5271311 |
| DBNDD2 | 7.51E-31 | -4.8378601 |
| DCAF8 | 1.03E-27 | -4.1124428 |
| DCTN4 | 1.17E-42 | -3.9372953 |
| DIS3L2 | 6.92E-30 | 3.89946868 |
| DOCK1 | 1.78E-56 | -4.9897879 |
| DYNC1LI1 | 5.12E-30 | -4.0480792 |
| EFCAB6 | 5.05E-25 | -3.6954477 |
| EIF1AD | 7.93E-125 | 6.37947139 |
| ERCC4 | 5.03E-09 | -4.3403268 |

| | | |
|---|---|---|
| F2RL2 | 2.25E-65 | -6.3013585 |
| GDI1 | 9.03E-54 | -3.9927621 |
| GEMIN4 | 2.30E-34 | 3.93722446 |
| GHRH | 1.56E-108 | -7.817523 |
| GPSM3 | 2.17E-37 | -3.9933201 |
| GTF2F2 | 4.29E-49 | 6.74322358 |
| GTF2H5 | 1.75E-70 | -6.1578766 |
| HNRNPA1 | 9.63E-17 | -3.5015788 |
| IFNGR1 | 5.15E-80 | -5.2430709 |
| IL17RD | 2.26E-31 | 4.50350572 |
| IL2RA | 4.79E-28 | -6.0764612 |
| IMPA1 | 3.39E-23 | 3.5340342 |
| INCA1 | 5.84E-57 | -5.5860732 |
| INHBC | 3.97E-92 | -4.2723328 |
| JUN | 7.05E-24 | -4.3067353 |
| KCNJ2 | 2.61E-79 | -4.2186756 |
| KIF1B | 1.72E-52 | -3.6607465 |
| LATS1 | 1.11E-35 | 4.08705167 |
| LINGO1 | 1.26E-15 | -4.697893 |
| LRP5 | 1.04E-36 | 3.66044924 |
| LTBP4 | 2.65E-41 | -3.6075486 |
| MADCAM1 | 3.42E-85 | -4.0795823 |
| MAP3K6 | 1.77E-66 | -3.9558842 |
| MAVS | 2.70E-44 | -3.7527404 |
| MCM6 | 1.31E-21 | -3.5672485 |
| MED25 | 1.46E-10 | -4.1479156 |
| MEP1A | 1.31E-52 | -3.8316063 |
| MITF | 4.97E-83 | -3.5094859 |
| MLLT1 | 1.65E-38 | -8.7694144 |
| MLLT6 | 5.72E-34 | -5.7953935 |
| MUSK | 3.83E-31 | -3.6255295 |
| NCR3 | 4.59E-71 | -4.6162681 |
| NDN | 1.18E-31 | -4.1169187 |
| NFX1 | 2.19E-39 | -7.1611357 |
| NOL3 | 7.73E-74 | -5.5187276 |
| NOL8 | 1.98E-07 | -3.901438 |
| NOSTRIN | 3.51E-18 | -4.5085776 |
| NR2F1 | 1.47E-46 | 4.41516163 |
| NR5A1 | 7.16E-56 | -5.4338555 |
| PAK1 | 1.32E-47 | -5.1234588 |
| PAM | 6.59E-13 | -4.2158121 |
| PDPN | 1.14E-63 | -5.7685641 |

| | | |
|---|---|---|
| PLIN3 | 3.89E-17 | -4.2807054 |
| PLK2 | 1.18E-28 | -4.4832599 |
| PMAIP1 | 1.57E-66 | 3.83360295 |
| PML | 5.49E-36 | -4.4926764 |
| POMP | 2.94E-12 | -3.6300316 |
| PPM1E | 2.33E-35 | -3.9880589 |
| PPP2R5E | 6.08E-14 | -3.7655002 |
| PPP3CC | 1.46E-34 | -4.3351609 |
| PRKG1 | 7.79E-50 | 4.41086621 |
| PTPRD | 3.00E-21 | -4.3537645 |
| RAB3GAP2 | 8.87E-50 | -4.199373 |
| RAP1GAP | 5.90E-52 | -7.1112586 |
| RBMXL2 | 2.20E-34 | -3.608101 |
| REXO4 | 4.04E-28 | -4.9149512 |
| RNF115 | 1.36E-30 | -5.2538633 |
| S100Z | 1.92E-32 | -5.6034288 |
| SAT1 | 8.61E-14 | -3.9051086 |
| SCG5 | 1.76E-58 | -3.7836285 |
| SEC24B | 4.41E-79 | -5.5362465 |
| SENP6 | 1.14E-44 | -4.4886736 |
| SERPINA10 | 1.34E-26 | -3.6655945 |
| SLC16A2 | 3.99E-21 | -3.7403086 |
| SLC37A1 | 4.49E-75 | -5.1199062 |
| SMARCA2 | 4.00E-64 | -3.6407725 |
| SNCAIP | 4.97E-21 | -3.5787914 |
| SNRNP200 | 1.65E-08 | -4.5538716 |
| SNX22 | 1.78E-24 | 3.76276335 |
| SOX2 | 8.15E-63 | 4.53096046 |
| SPRR2A | 8.60E-72 | -4.2878226 |
| ST13 | 1.79E-39 | -5.3182038 |
| SUMF2 | 1.27E-50 | -3.541777 |
| SYN1 | 2.80E-13 | -5.0970774 |
| TEP1 | 1.98E-38 | -5.4549138 |
| TP53RK | 8.31E-41 | 3.53965215 |
| TPP1 | 1.98E-68 | -4.498507 |
| ULBP3 | 1.02E-18 | -4.6317657 |
| VSNL1 | 1.94E-27 | -5.3289795 |
| WEE1 | 1.50E-28 | 3.80049792 |

**Supplementary methods**

***Curvature-based edge ranking analysis***

Using ORC, the strength of connections originating from a gene can be summarized by computing the scalar curvature for a given node. Scalar curvature is the summation of edge curvature values originating in the node (gene) of interest and is formally defined below.

$$k_i = \sum_{j \sim i} k_{i,j}$$

ORC for RNA-Seq and CNA data was computed separately for each patient. To consolidate the geometric information induced by the nodal weights (RNA-Seq and CNA), we computed the pure topological curvature (wherein all the weights were set to be 1) and subtracted that from each of the original curvature values for all patients. Once computed, gene pairs were then ranked based on the computed value which we call the strength of curvature.
